# Warm blood meal increases digestion rate and milk protein production to maximize reproductive output for the tsetse fly, *Glossina morsitans*

**DOI:** 10.1101/2022.08.07.501667

**Authors:** Joshua B. Benoit, Chloé Lahondère, Geoffrey M. Attardo, Veronika Michalkova, Kennan Oyen, Yanyu Xiao, Serap Aksoy

## Abstract

The ingestion of blood represents a significant pressure that immediately increases water, oxidative, and thermal stress, but provides a significant nutrient source to generate resources necessary for the development of progeny. Thermal stress has been assumed to solely be a negative byproduct that has to be alleviated to prevent stress. Here, we examined if the short thermal bouts incurred during a warm blood meal are beneficial to reproduction. To do so, we examined the duration of pregnancy and milk gland protein expression in the tsetse fly, *Glossina morsitans*, that consumed a warm or cool blood meal. We noted that an optimal temperature for blood ingestion yielded a reduction in the duration of pregnancy. This decline in the duration of pregnancy is due to increased rate of blood digestion when consuming warm blood. This increased digestion likely provided more energy that leads to increased expression of transcript for milk-associated proteins. The shorter duration of pregnancy is predicted to yield an increase in population growth compared to those that consume cool or above host temperatures. These studies provide evidence that consumption of a warm blood meal is likely beneficial for specific aspects of vector biology.

## Introduction

Tsetse flies are important vectors of disease caused by African trypanosomes, known as sleeping sickness in humans and Nagana in animals. An effective way to combat this disease involves vector control applications, which are highly efficient due to the low reproductive output of tsetse. Tsetse flies are one of the few insects that employ viviparity (Benoit et al. 2015; Meier, Kotrba, and Ferrar 1999), characterized by the provision of nutrients beyond egg yolk to support embryonic development and growth of larva to full term within the female uterus. Female tsetse flies produce a single mature third instar larva during each gonotrophic cycle following a 4-6 day period of intrauterine gestation (Benoit et al. 2015; Tobe 1978). Thus, these K-strategists produce only a modest 8-10 progeny per female, per lifetime (Benoit et al. 2015; Tobe 1978; Lord et al. 2021; Michalkova et al. 2014). A critical adaptation underlying this reproductive strategy is the modification of the female accessory gland to secrete milk into the uterus for larval consumption. The milk is composed of proteins and lipids emulsified in an aqueous base (Cmelik, Bursell, and Slack 1969; Benoit, Attardo, et al. 2014). During lactation, at least 6-10 mg of nutrients dissolved in 12-14 mg of water are transferred to the larva. Molecular characterization of tsetse milk revealed 12 major milk gland proteins, including Transferrin (Guz et al. 2007; Benoit, Attardo, et al. 2014), a Lipocalin (Milk Gland Protein1, MGP1 (Attardo et al. 2006; Benoit, Attardo, et al. 2014), nine tsetse-specific milk proteins (MGP2-10; (Attardo et al. 2019; International Glossina Genome Initiative 2014; Benoit, Attardo, et al. 2014)), and Acid Sphingomyelinase 1 (aSMase1; (Benoit et al. 2012)). Transcriptomic analysis revealed that the milk proteins represent over 47% of the total transcriptional output in lactating flies. The contribution of the milk protein transcripts declines to less than 2% of the total output two days after the lactation cycle, at parturition (Benoit, Attardo, et al. 2014). This high investment in the progeny indicates that nutrients need to be rapidly processed for storage or direct allocation to the developing larvae. As an obligate blood feeding insect, all nutrients are derived from the consumption of a blood meal.

Previous studies have shown that the ingestion of a warm blood meal can be detrimental to blood feeding insects, where an increase in from ambient (24-26°C) to the temperature of the host (37-40°C) can cause thermal damage (Lahondère and Lazzari 2012; Benoit et al. 2011; Lahondère et al. 2017; Benoit et al. 2019). Tsetse flies are not immune to this process, as a drastic increase in body temperature occurs during blood feeding (Lahondère and Lazzari 2015). Importantly, as increased temperature is critical for the digestion process (McCue et al. 2016), it could be important that a warm blood meal may allow for more rapid processing of the blood meal. This could subsequently allow for increased reproductive output. To do so, we examined the impact of feeding on blood at various temperatures on pupal mass and duration of pregnancy. This was followed by the measuring of blood digestion and the expression of milk protein along with population modeling. These results indicate that the consumption of a warm blood meal is likely critical for tsetse flies to reach their maximum fecundity.

## Materials and Methods

### Flies

*Glossina morsitans morsitans* (Westwood, 1851) were reared at Yale University. Flies were maintained on bovine blood meals provided through an artificial feeding system at 48h intervals (Moloo 1971) that can be varied in temperature. Flies were maintained under 12h:12h light:dark at 25°C.

### Thermal imaging and contact thermal changes during blood meal

Thermal imaging was conducted as in Lahondère and Lazzari (2015). Briefly, a thermographic camera (PYROVIEW 380L compact, DIAS infrared GmbH, Germany; spectral band: 8–14 mm, uncooled detector 2D, 384×288 pixels) with a macro lens (pixel size 80 µm; *A*=60 mm; 30°×23°) was used for data acquisition (Lahondère and Lazzari 2015). The emissivity was set at 0.98 as determined previously for insect cuticle (Stabentheiner 1987).

Direct temperature changes were monitored with the use of a thermocouple (Omega) attached to the tsetse fly using petrolatum on the top of the abdomen, and tsetse flies were placed on an artificial feeding system as described previously. Temperatures of eight females were monitored during feeding with a HHM290 thermometer (Omega). All statistics were conducted with the use of R packages (3.6.3).

### Blood digestion

Blood digestion quantification was determined according to previous methods (Langley et al. 1978). Briefly, four and sixteen hours following a blood meal, the digestive system was removed from the proventriculus to the point of Malpighian tubules insertion. All fat body and tracheal tubes were removed. If the gut ruptured, the sample was not used in further analyses. The guts and their contents were dried at 50°C in the presence of Drierite (Xenia, OH) until the mass was constant. The dry mass was set as the relative amount per 50 mg of blood to allow comparison to previous studies (Langley et al. 1978). All statistics were conducted with the use of R packages (3.6.3).

### RNA and protein isolation

RNA and protein isolations were performed using Trizol reagent on whole flies, and isolated milk gland/fat body samples were isolated following the instructions provided by the manufacturer (Invitrogen). RNA was cleaned with an RNeasy Mini Kit (Qiagen). Complementary DNA was synthesized using a Superscript III reverse transcriptase (Invitrogen) kit from 1µg of the total RNA isolated from each sample.

### Quantitative PCR

Transcript levels for *mgp1, asmase1, mgp7*, and *trf* (gene sequences acquired from the *Glossina* genome project, vectrobase.org) were determined via qPCR by employing the CFX real-time PCR detection system (Bio-Rad, Hercules, CA) with primers specific to each target gene. Primers were used in previous studies (Benoit, Attardo, et al. 2014; Benoit et al. 2012, 2018). All readings were obtained on four biological replicates that were normalized to tsetse *tubulin* expression levels. CFX Manager software version 3.1 (Biorad) was used to quantify transcript expression of each gene and conducted according to methods developed in previous studies (Benoit et al. 2012; Benoit, Hansen, et al. 2014). All statistics were conducted with the use of R packages (3.6.3).

### Population modeling

We developed a mathematical model to examine the population dynamics of tsetse flies using Matlab (Mathworks). We consider three compartments representing immature female (P), matured female within the first deposition period (A_1_), and matured female in other deposition periods (A_2_,). Here, the immature class is assumed to include both populations of larvae and pupae. We let 1/*d*_*p*_ and 1/*d*_*a*_ be the expected duration of the immature and matured females. *a*_1_ and *a*_2_ are the development rates from *P* to *A*_1_ and *A*_1_ to *A*_2_, respectively. Our model then is followed as

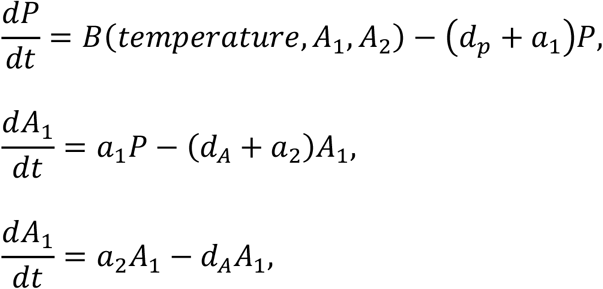

where *B*(*temperature, A*_1_, *A*_2_) represents the temperature dependent birth rate. Based on the analysis in Hu et al. 2008 and Benoit t al. 2018, we calculated the average expected number of female offspring of a female parent per unit time during each deposition as

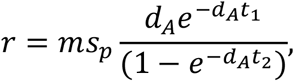

where *m* is the proportion of offspring that are female; *s*_*p*_ is the survival rate of deposited larva; *t*_1_ and *t*_2_ are the duration of the previous and current deposition cycle.

During the first deposition, to calculate the average daily birth rate of the group *A*_1_, we have *t*_1_ = *t*_2_ + 8 as the duration of the first deposition is longer than that of other deposition cycles. To estimate the populations *Glossina* after 1 year, with the initial population to be 100 pupae and 100 adults, we calculated the average of the populations for 100 runs of the model simulations under different temperature settings. For each temperature setting and each run, we use rando-sampled life span for adult populations, and the durations of each deposition till the end of 1-year-period are also randomly sampled from our experiment dataset.

## Results

### Temperature changes in tsetse flies associated with a blood meal

Thermal imaging revealed that the flies’ body temperature increased to near the temperature of the ingested blood during feeding (37°C) (Fig. 1A). When the temperature was tracked with a probe when feeding on warm or cool blood, there was a much greater increase in temperature of the fly thorax when feeding on warm (37-38°C) rather than cool blood (30-31°C) (Fig. 1B). These results indicate that a warm blood meal increases the flies’ body temperature more drastically than during the ingestion of a cool blood meal.

**Fig. 1.**
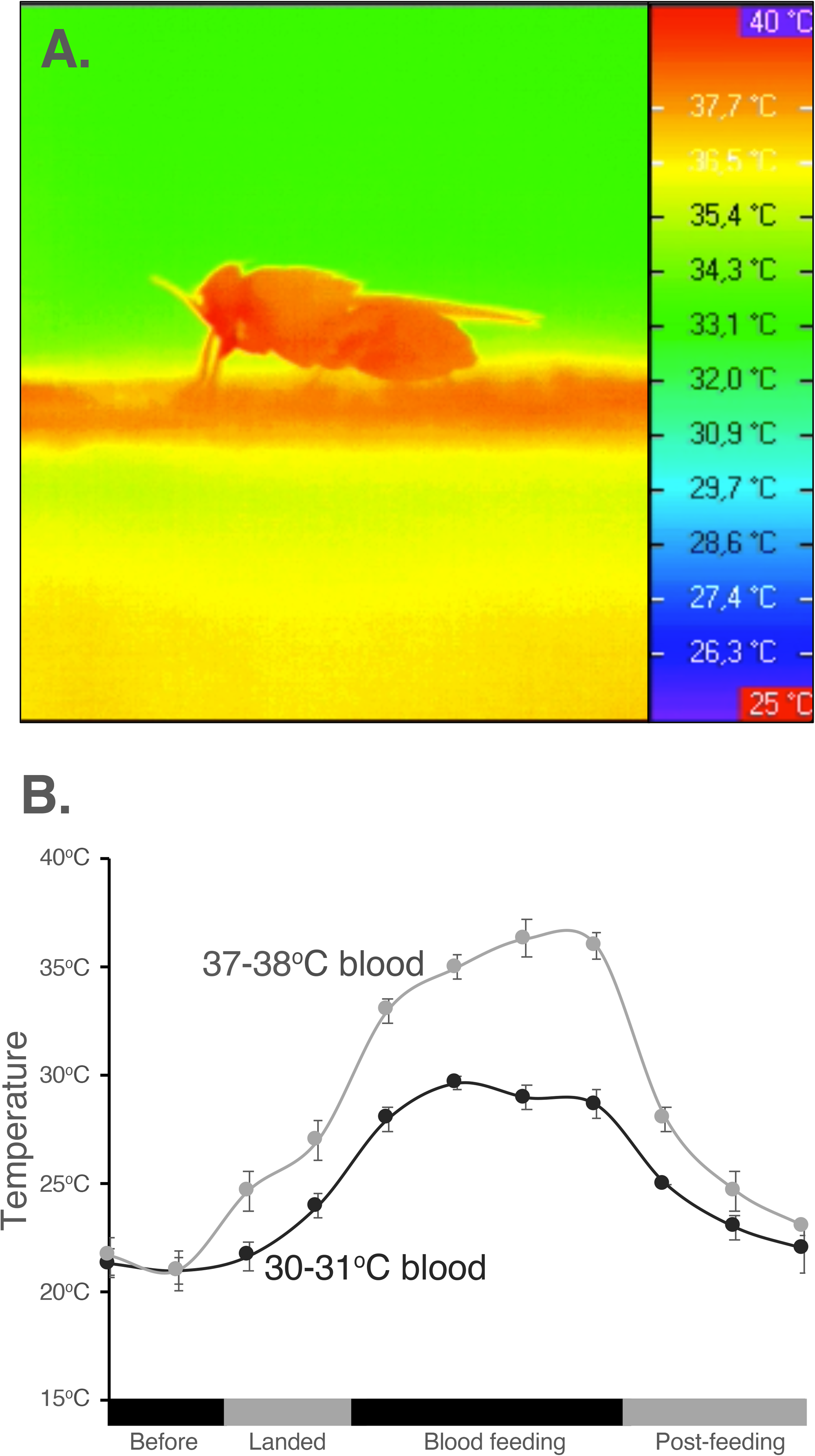
Thermal changes in tsetse flies during the consumption of blood. A. Thermal image of tsetse fly when feeding on warm blood. B. Thermal changes in tsetse fly abdomen when consuming warm and cool blood. There is a significant reduction in temperature from when the fly landed on the host until after the completion of feeding.

### Pupae size and duration of pregnancy

When a blood meal was offered at multiple temperatures, no difference in the size of the pupae produced was found (Fig. 2A). This is not surprising as few differences are noted in the size of pupae produced when larviposition occurs. When the duration of pregnancy was assessed, there was a significant reduction in the duration of pregnancy when flies were fed at 38°C (Fig. 2B). This reduction only occurred at this temperature and no longer occurred when the blood was at 41°C. This indicated that a warm, but not hot, blood meal may be beneficial to tsetse flies by reducing pregnancy duration.

**Fig. 2.**
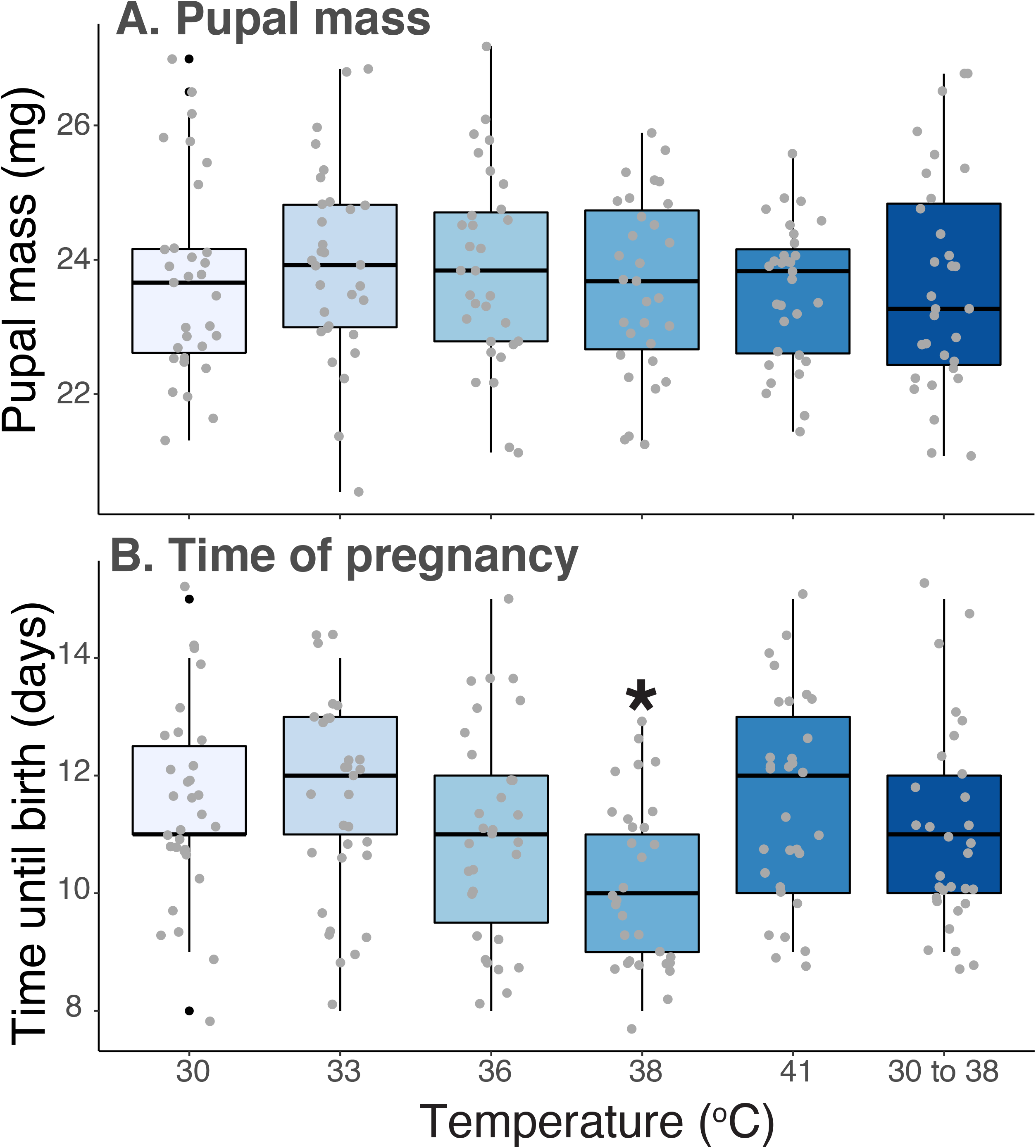
Progeny size and pregnancy duration changes in relation to blood meal temperature. A. Pupal mass and B. time of pregnancy in relation to the composition of blood at multiple temperatures. Statistical differences were assessed with the use of ANOVA followed by post-hoc tests.

### Blood processing and milk protein synthesis

When the mass of the midgut was examined as a proxy for digestion, we noted that there was a delay in the processing of the blood meal (Fig. 3) in flies that consumed a cool blood meal. After twelve hours, the dry mass of these flies was significantly higher than the individuals that consumed a warmer blood meal. The expression of milk gland protein genes was lower in the flies that were offered a cool blood meal twice before 6 days into a pregnancy cycle. The results suggest that ingesting a cool blood meal leads to a delay in the pregnancy cycle, likely due to a slower blood digestion and a delay/reduction in the production of milk proteins.

**Fig. 3.**
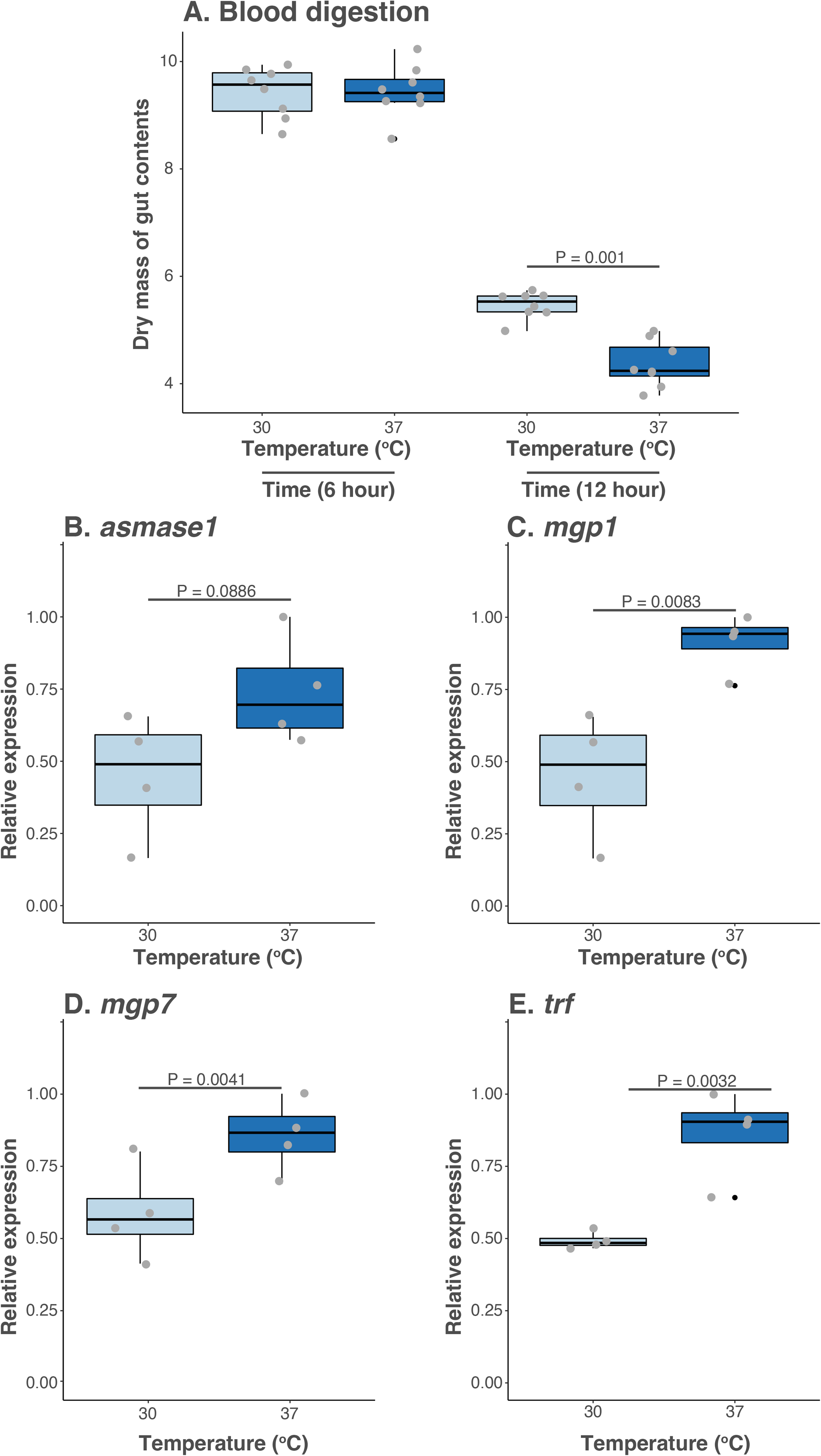
Blood digestion and milk gland protein transcript abundance. A. Dry mass of the *G. morsitans* midgut 6 and 12 hours following a blood meal. B-E. Transcript levels for four milk gland protein when flies consumed cool (30°C) or warm blood (37=8°C). Statistical differences were assessed with the use of a Student *t*-test.

### Population modeling

To determine the impact of warm or cold blood meals on the population dynamics of *Glossina*, we used the model generated in this study to estimate the populations of tsetse flies after 1 year with the initial population consisting of 10 pupae and 10 adults within their first deposition period. These parameters were based on previous studies (Hu et al. 2008; Benoit et al. 2018). We evaluated the average of the populations for 100 runs of the model simulations under different temperature settings. We determined that the average total population when feeding on warm blood (37-38°C) after 1 year is significantly higher than those fed on blood at other temperatures even with small initial population (Fig. 4A), but if individuals fed on a warm blood meal and then cool this benefit was eliminated (Fig. 4B).

**Fig. 4.**
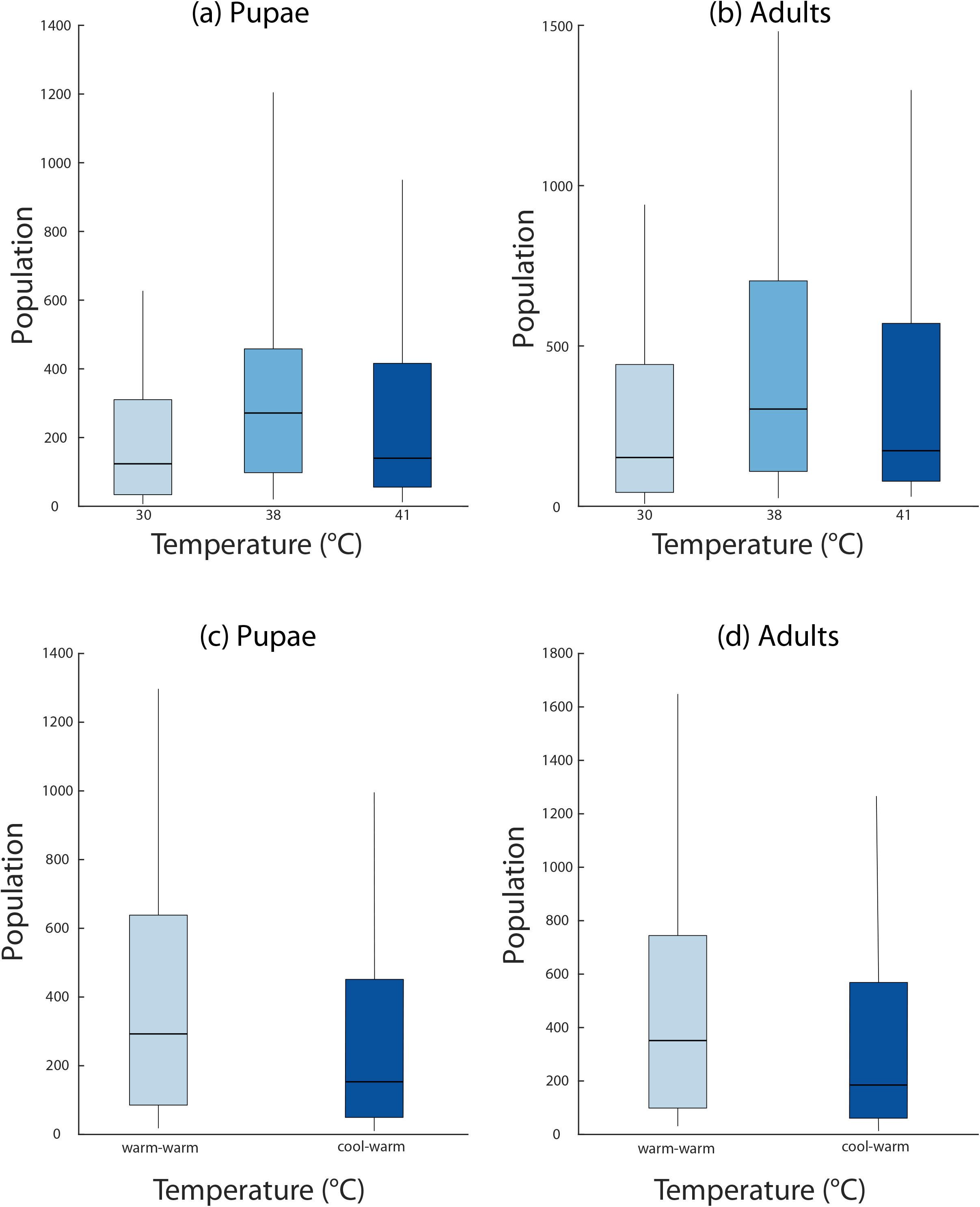
Population modeling based on changes in pregnancy duration following feeding. A and B. Pupae and adult population levels when consuming cool, warm, and hot blood. Values for all temperatures are provided in Figure S1. C. A cool blood meal (30°C) followed by a second warm blood meal (38°C) each week still leads to a lower population growth than those offered a warm blood meal (38°C) for all meals.

## Discussion

In this study, we show that the temperature of a blood meal can impact the productive output of tsetse flies by reducing the overall duration of pregnancy. This decreased pregnancy cycle is likely due to an increased blood meal processing, which yields a decreased production of milk proteins. This reduced pregnancy cycle likely yields a higher population growth when compared to flies that consumed either cool blood or blood at temperatures above that of a normal host. These results suggest that ingestion of warm blood is critical to tsetse fly reproduction.

Blood ingestion represents a period of extreme stress that includes increased oxidative stress (Sterkel et al. 2017), toxicity from the overabundance of specific amino acids and ions, and an overabundance of water and iron products. Blood feeding insects tolerate these stresses via a combination of adaptations developed over the course of their evolution into this life strategy. Specifically, several mechanisms allow insects and other blood feeding arthropods to remain cool during a blood meal. This includes a counter-current heat exchange that allows kissing bugs to remain cool while feeding (Lahondère et al. 2017) or evaporative cooling through urination that has been documented in mosquitoes and ticks (Benoit et al. 2019; Lahondère and Lazzari 2012). Damage prevention at the cellular level is accomplished through the expression of heat shock proteins (Benoit et al. 2011, 2019; Lahondère et al. 2017). Suppression of these proteins impacts the processing of the blood and egg production. Thermal stress has been mainly assumed to be a detriment during blood feeding(Lahondère et al. 2017; Benoit et al. 2019), but as digestion is directly tied to temperature in tsetse flies, a warm bloodmeal may be critical to digestion (McCue et al. 2016). Based upon our results, digestion is delayed when cool blood is consumed, suggesting that increased temperature during the blood meal is critical for tsetse flies to rapidly process their blood meal.

The rapid processing and incorporation of the blood meal is critical for tsetse flies to reach their maximum reproductive output. Immediately after larval deposition, there is a rapid period of milk gland involution (Benoit et al. 2018), which allows the female to begin to accumulate nutrient reserves. During this time, flies consume a fewer number of blood meals, usually only two-three, that function directly as a source of energy and build the lipid reserves (Attardo et al. 2012; Langley et al. 1981; Benoit et al. 2015). Likely, the delay in digestion we observed leads to a delay in the production of milk proteins as resources from the blood meal are not yet available. Importantly, the consumption of blood at suboptimal temperatures leads to a slight reduction in population growth, suggesting that tsetse fly maximum reproductive output requires feeding on warmer blood.

In conclusion, we provide evidence that consumption of warm blood meal may not only be detrimental to blood feeding insects as previously observed in other systems. Rather the impact is species specific and can be nuanced, as specific aspects may be beneficial for the insect physiology, such as the processing of the blood meal and reproductive output in tsetse flies.

## Acknowledgements

We thank Yineng Wu for her technical expertise. Funding was provided by the National Institutes of Health (AI081774 and F32AI093023) and International Atomic Energy Agency (Technical Contract No. 17713/R0). We thank Claudio Lazzari for the use of thermal images generated in his research lab.

## Figure legends

**Figure S1.**
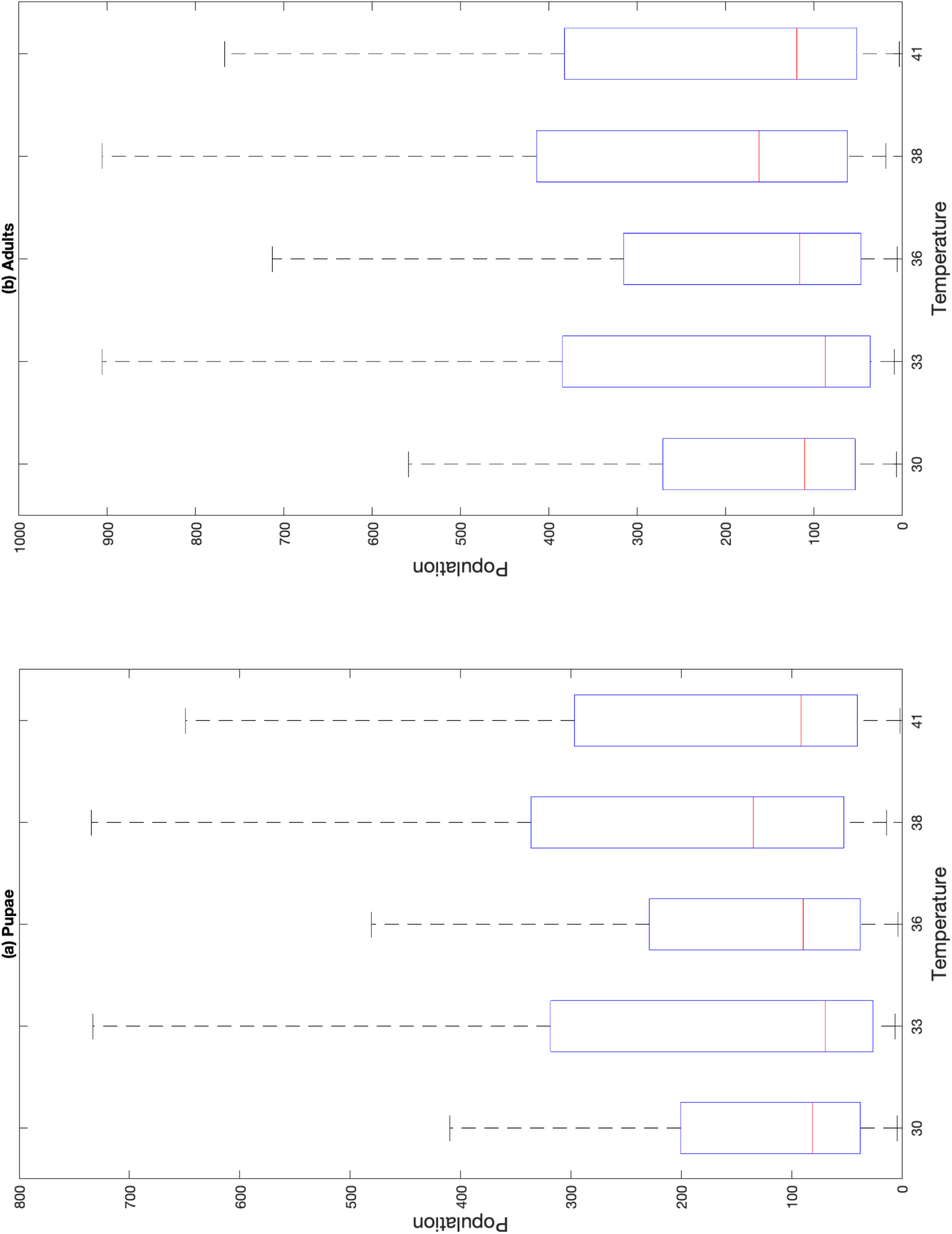
Full population modeling. Population growth rates observed when tsetse flies consumed blood at different temperatures.

